# Antiviral activity of Molnupiravir precursor NHC against SARS-CoV-2 Variants of Concern (VOCs) and implications for the therapeutic window and resistance

**DOI:** 10.1101/2021.11.23.469695

**Authors:** Tessa Prince, I’ah Donovan-Banfield, Hannah Goldswain, Rebekah Penrice-Randal, Catherine Hartley, Saye Khoo, Tom Fletcher, Julian A. Hiscox

## Abstract

Several regulatory agencies have either licensed or given emergency use approval for treatment of patients at risk of developing severe COVID-19 with the anti-viral drug, Molnupiravir. Recent trials involving Molnupiravir suggested the drug was not as efficacious as earlier studies suggested. This study aimed to: (i) determine the effectiveness of the Molnupiravir active metabolite (NHC) against different SARS-CoV-2 Variants of Concern (VoCs), (ii) establish the therapeutic window of NHC in a human lung cell model, and (iii) and evaluate the genetic barrier to resistance. Dose response assays were performed in parallel to determine the IC50 (the concentration required to inhibit virus titre by 50%) of NHC against different variants. Human ACE-2 A549 cells were treated with NHC at different time points either before, during or after infection with SARS-CoV-2. Multiple passaging in the presence or absence of drug was used to evaluate whether resistance occurred. To obtain genomic information, virus was sequenced at regular intervals. After 20 passages in the presence of the drug, dose response assays and sequencing showed the virus did not appear to have developed resistance. The drug had equivalent activity against four VOCs ranging from 0.04 to 0.16μM IC50. The efficacy of the drug diminished when applied after 24 hours post-infection. Our results suggest that earlier administration in patients, perhaps pre- or post-exposure rather than symptom onset, would be a more effective treatment option.

## 1.0 Introduction

SARS-CoV-2 emerged in China in late 2019 and is the causative agent of the disease known as COVID-19. While vaccination efforts have been largely successful in preventing severe disease, many people worldwide remain unvaccinated. Additionally, important groups of patients, such as those on immunosuppressive therapies, mount suboptimal responses to vaccines (Kearns et al., 2021). For people in these categories, initially successful alternative countermeasures such as monoclonal antibodies and protease inhibitors are declining in their effectiveness as the virus evolves (Cox et al., 2022; Moghadasi et al., 2022). Therefore, treatments that have longevity in the face of virus evolution are required.

Anti-virals can target either specific viral proteins or act non-specifically across the viral genome. Three anti-virals for SARS-CoV-2 have been licensed in the UK: Nirmatrelvir/Ritonavir (Paxlovid), Remdesivir and Molnupiravir (Haddad et al., 2022), each with their own advantages and disadvantages.

Paxlovid is an oral combination therapy of protease inhibitors targeting the SARS-CoV-2 main protease (M^pro^). It has been successfully used clinically, though must be used with care due to its drug-drug interactions. However, there are some *in-vitro* indications that resistance can develop to Nirmatrelvir (Jochmans et al., 2023; Zhou et al., 2022). The nucleoside analogue Remdesivir incorporates into the viral RNA strand, and causes chain termination, directly inhibiting the proteins involved in RNA synthesis (Kokic et al., 2021). However, the drug must be given intra-venously, requiring a visit to hospital. Additionally, resistance to the drug has also been shown to develop *in vitro* (Szemiel et al., 2021). Finally, another nucleoside analogue, which acts to indirectly inhibit SARS-CoV-2 replication through the induction of transition mutations (e.g. C -> U or G -> A changes) is Molnupiravir (Kabinger et al., 2021). This results in error catastrophe during viral replication.

Molnupiravir is an orally bioavailable pro-drug that is converted enzymatically *in-vivo* to its active form ß-D-N4-hydroxycytidine (NHC). Molnupiravir treatment reduces viral load and block transmission in several animal models (Abdelnabi et al., 2021; Cox et al., 2021; Wahl et al., 2021).

Molnupiravir has been evaluated in several large clinical trials for the treatment of COVID-19. The MOVe-OUT trial held in the USA suggested a subtle effect of the drug with an estimate of reduction in risk of hospitalisation or death from 9.7% to 6.8% in unvaccinated individuals (about 30% reduction). In this trial 47.7% received the drug within 3 days or less from symptom onset, the rest within 5 days. Molnupiravir administration resulted in faster normalisation of clinical indicators compared to placebo, and those who received the drug earlier had larger reductions in viral load compared to baseline (Jayk Bernal et al., 2022). A post-hoc analysis of immunocompromised patients in the MOVe-OUT trial further found Molnupiravir reduced incidence of hospitalisation or death (Johnson et al., 2023). The open-label PANORAMIC trial from the UK has recently suggested that Molnupiravir, when given to vaccinated individuals at a median of three days from symptom onset, did not significantly reduce risk or hospitalisation/death, but did result in a faster self-reported time to recovery and reduced viral load (Butler et al., 2022). The AGILE trial in the UK supported this result suggesting a shorter time to PCR negativity and some moderate anti-viral activity in both vaccinated and unvaccinated individuals compared to placebo, though the evidence was not conclusive (Khoo et al., 2022). Finally, another Phase 2a double-blind, placebo controlled, multicentre trial evaluated the safety and tolerability of the drug and found that 92.5% of Molnupiravir treated patients achieved viral RNA clearance (measured by PCR) compared to 80.3% who received placebo. Likewise, isolation of live virus decreased after administration of the drug (Fischer et al., 2021).

The development and assessment of anti-viral drugs is a notoriously long process. An anti-viral drug should be effective against both current and future variants. Additionally, timing and dosage of treatment should be optimised to reduce the development of resistance. There have been varied reports for the efficacy of Molnupiravir in mixed target populations. Therefore, we have further characterised Molnupiravir by evaluating its activity in two different cell lines and identifying if delays in treatment lead to sub-optimal activity. Furthermore, we evaluated the effect of repeated exposure of SARS-CoV-2 to Molnupiravir.

## 2.0 Materials and Methods

### 2.1 Compound

NHC (Alsachim) was supplied as a 1mg powder (purity 95%) and was resuspended in 1ml DMSO to provide a 4.07mM stock solution. This was diluted in viral maintenance media (DMEM containing 2% FBS and 0.05mg/ml gentamicin) for experiments using a range of concentrations.

### 2.2 Cell culture

Human ACE2-A549 (hACE2-A549 were the kind gift of Oliver Schwartz (Buchrieser et al., 2020). These were cultured in DMEM with 10% FBS and 0.05mg/ml gentamicin with the addition of 10μg/ml Blasticidin (Invitrogen). Only passage 3-10 cultures were used for experiments. Calu-3 cells (HTB-55; ATCC) were cultured in DMEM with 10% FBS and 0.05mg/ml gentamicin. Vero/hSLAM cells (PHE) were grown in DMEM with 10% FBS and 0.05mg/ml gentamicin (Merck) with the addition of 0.4mg/ml Geneticin (G418; Thermofisher) at 37°C/5% CO_2_.

### 2.3 Viral Culture

Virus stocks were grown in Vero/hSLAM cells using DMEM containing 2% FBS, 0.05mg/ml gentamicin and 0.4mg/ml geneticin and harvested 72 hours post inoculation. Virus stocks were aliquoted and stored at −80°C. The titre of stocks (PFU/ml) was determined by plaque assay. RNA from viral stocks were sequenced by Oxford Nanopore long read length sequencing on flow cells run on GridION.

### 2.4 Sequencing of viral stocks

Sequencing libraries for ARTIC amplicons were prepared following the ‘PCR tiling of SARS-CoV-2 virus with Native Barcoding’ protocol provided by Oxford Nanopore Technologies using LSK109 and EXP-NBD196. The artic-ncov2019 pipeline v1.2.1 (https://artic.network/ncov-2019/ncov2019-bioinformatics-sop.html) was used to filter the passed fastq files produced by Nanopore sequencing with read lengths between 400 nt and 700 nt for ARTIC amplicons. This pipeline was then used to map the filtered reads on the reference SARS-CoV-2 genome (MN908947.3) by minimap2 and assigned each read alignment to a derived amplicon and excluded primer sequences based on the ARTIC V3 and V4 primer schemes in the bam files. Primer-trimmed bam files were further analysed using DiversiTools (http://josephhughes.github.io/DiversiTools/) with the “-orfs” function to generate the ratio of amino acid change in the reads and coverage at each site of protein in comparison to the reference SARS-CoV-2 genome (MN908947.3). The amino acids with highest ratio and coverage > 10 were used to assemble the consensus protein sequences. Pangolin (https://pangolin.cog-uk.io/) was used to confirm lineages of each viral stock used in experiments.

### 2.5 *In-vitro* cytotoxicity of NHC

Human ACE2-A549 cells and Calu-3 cells were plated at 2 x 10^4^ cells per well in a clear bottomed white 96 well plate. Twenty-four hours later the medium was replaced with media containing NHC at different concentrations. At 72 hours post-exposure, cell viability was measured by CellTiter-Glo assay (Promega) as per the manufacturer’s instructions.

### 2.5 Anti-viral activity of NHC against SARS-CoV-2 Variants of Concern (VoCs)

Human ACE2-A549 cells and Calu-3 cells were grown to confluency and infected at an MOI (Multiplicity of Infection) of 0.1 in either DMEM with 2% FBS and 0.05mg/ml gentamicin, or in the same media containing 0.01μM, 0.1μM, 1μM or 10μM NHC by allowing virus to adsorb to cells in a volume of 100μl for one hour at 37°C, and then topping up to 500μl with the relevant media afterwards. A mock infected control and a DMSO control were included in each experiment and experiments were repeated a minimum of 3 times. After 72 hours, supernatants were collected and stored at −80°C until viral titre was determined by plaque assay. The inhibitory potency of NHC measured as the absolute IC50 was defined as the concentration of drug that resulted in a 50% reduction in the number of plaques compared to untreated controls.

### 2.6 Pre-exposure and Post-exposure to NHC

For pre-exposure experiments, media was removed from hACE2-A549 cells and replaced with media containing NHC two hours prior to infection. This was then removed for infection, which was performed as described above. For concomitant and post-exposure experiments, infections were performed in media as described above. At two, four, 24 or 48 hours post-infection and concomitant infection media was removed from cells. The cells were washed twice with PBS, and the media replaced with media containing 0μM, 0.01μM, 0.1μM, 1μM or 10μM NHC NHC. After 72 hours post-infection, supernatants were collected and stored at −80°C until viral titre was determined by plaque assay.

### 2.7 Serial Passage experiments for detection of resistance

To determine if Molnupiravir resistance would occur through repeated exposure to low doses of the drug, SARS-CoV-2 was passaged every 2-3 days in Calu-3 cells in either the presence or absence of the approximate IC50 of the drug (0.1μM). Passaging was done by taking 100μl of the supernatant and adding it to fresh cells for one hour at 37°C before topping up with the relevant media. At passages 1, 5, 15 and 20 the supernatant was collected, titred by plaque assay, and (i) used to perform dose response assays to the drug (as previously described) at an MOI of 0.01 for each passage and (ii) RNA extracted and NimaGen-illumina sequencing performed. An illustration of the experimental outline if provided in figure 1.

**Figure 1.**
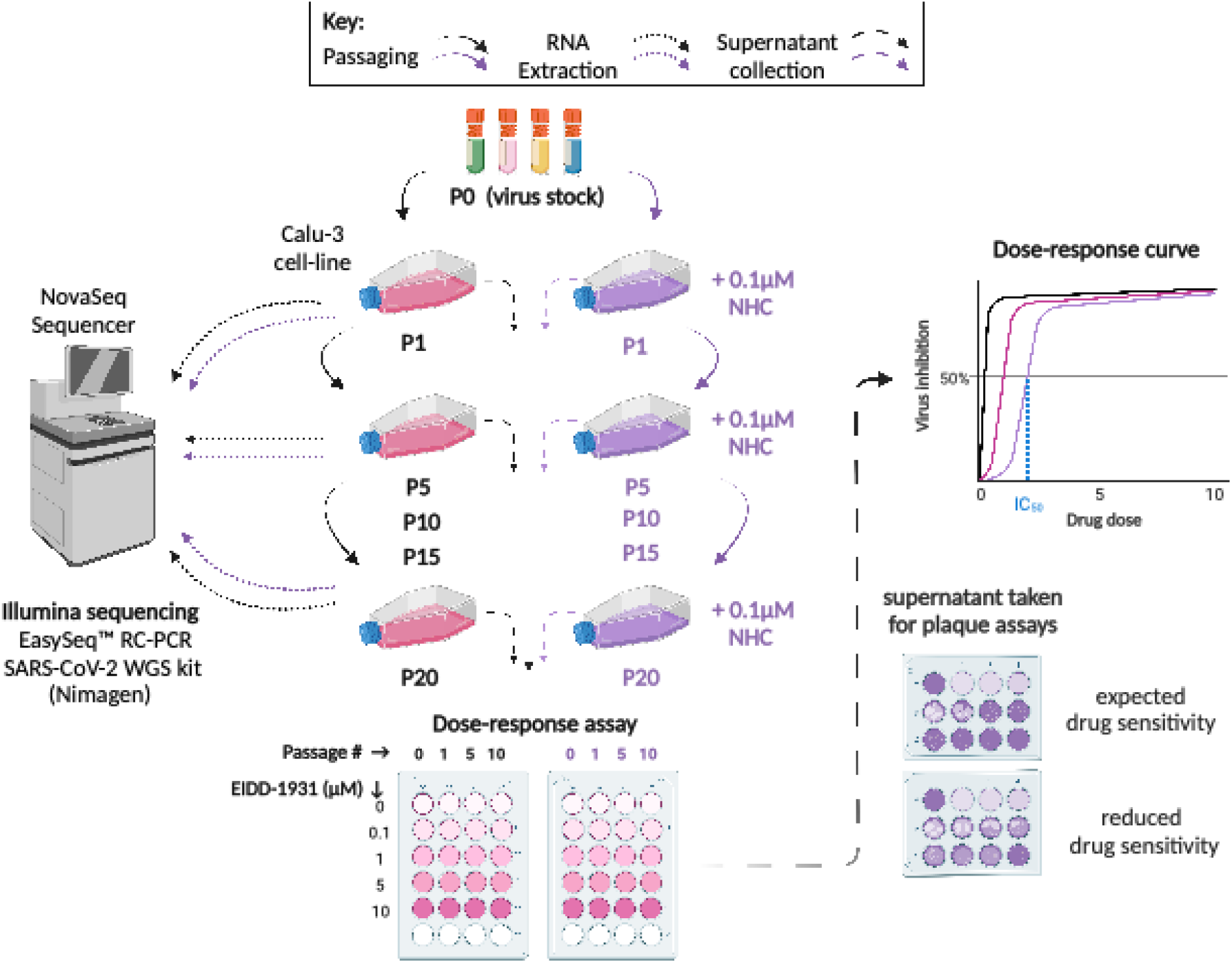
Experimental outline for measurement of resistance of SARS-CoV-2 to NHC. Calu-3 cells were infected with SARS-CoV-2 at an MOI of 0.01 in either the presence or absence of the approximate IC50 of the drug (0.1μM) and then passaged every 2-3 days. Virus was passaged by taking 100μl of the supernatant and adding it to fresh cells for one hour at 37 °C before topping up with the relevant media with or without NHC. At passages 1, 5, 15 and 20 the supernatant was collected, titred by plaque assay, and (i) used to perform dose response assays to the drug (as previously described) at an MOI of 0.01 for each passage and (ii) RNA extracted and NimaGen-illumina sequencing performed.

### 2.8 NimaGen-Illumina Sequencing and Bioinformatic analysis

NimaGen libraries were prepared and amplified as described previously before being sequenced on the NovaSeq 6000 machine. Bioinformatic analysis was likewise performed as previously described (Donovan-Banfield et al., 2022). Briefly, the raw sequencing data was processed using two different pipelines. The first pipeline, EasySeq_covid19 (version 0.9, code available at https://github.com/JordyCoolen/easyseq_covid19), performs quality control steps, maps to the reference genome (Wuhan-Hu-1; NC045512.2 https://www.ncbi.nlm.nih.gov/nuccore/NC_045512.2), variant calls and generates a consensus genome for each sample (Coolen et al., 2021). Default parameters were used and are as follows: variant call threshold = 0.5; variant calling quality threshold = 20; variant calling minimum depth = 10. Pangolin (version 4.0.6) was used to assign SARS-CoV-2 lineage, with maximum ambiguity set at 0.31 (https://pangolin.cog-uk.io/) (O’Toole et al., 2021). The second method, DiversiTools (code available at https://github.com/josephhughes/DiversiTools), uses the primer-trimmed alignment file (named as [sampleID]_L001.final.bam) and its associated index file (produced in the EasySeq pipeline) along with the reference genome and a coding region file to analyse the minor genomic variation and predict the amino acid sequence based on the genomic data. DiversiTools allows for an in-depth analysis of viral diversity in each sample, rather than just the consensus/dominant genomic information. Data visualisation was conducted in R (version 4.0.2), using the tidyverse package (version 1.3.2) for data manipulation. All plots were created using ggplot2 (version 3.3.6). Figures were compiled using cowplot (version 1.1.1) and magick (version 2.7.3) packages. Data and code are available at https://doi.org/10.5281/zenodo.7594012.

### 2.9 Statistical analysis

A one-way ANOVA with post-hoc Tukey test was used to evaluate the *in-vitro* cytotoxicity data. For dose response curves, the absolute IC50 values were calculated using GraphPad Prism 9.5, using a non-linear 4-parameter logistic regression. An extra sum-of-squares F-test was used to determine if there was any evidence that that the data sets in each logistic regression differed from each other. T-tests comparing the IC50 values for hACE2-A549 cells and Calu-3 cells were used to determine if the IC50 values differed between cell lines. As an additional analysis, a one-way ANOVA was performed on the recorded IC50 values for the serial passaging experiment. P was considered significant if <0.05 and a minimum of three replicates was performed for each experiment. All plots were created using GraphPad Prism 9.5 and figure 1 was created using Biorender.com.

### 3.0 Results and Discussion

To find the appropriate non-toxic concentration range of NHC in hACE2-A549 and Calu-3 cells, Cell-titer Glo Assays (Promega) were used to measure % ATP production in cells. These were treated with DMSO or different concentrations of NHC diluted in culture medium compared to mock untreated cells. There was a slight increase (105.9%) in the ATP production in cells at the lowest concentration of drug, 0.01μM (p=0.02) in hACE2-A549 cells. The only concentration of drug to inhibit the ATP production was 10μM for both cell lines (89.74±1.03%, and 81.2 ± 1.6%, p>0.0001) (Supplementary Figure 1a and b). However, cell monolayers appeared completely intact, and therefore this dose was used as the upper limit in the dose response assays.

### 3.1 NHC has equivalent efficiency against different VOCs *in vitro*

NHC has been evaluated against different VOCs in various cell lines, with its IC50 (the concentration of drug required to reduce the amount of live virus by 50%) reported between 0.08-10μM (Bojkova et al., 2022; Lee et al., 2021; Rosenke et al., 2021; Sheahan et al., 2020; Stegmann et al., 2021; Zhou et al., 2021). Therefore, before we could determine the treatment window, we needed to determine what the IC50 was in our cell lines and if it was equivalent for different VOCs. Molnupiravir, as a nucleoside analogue, acts by mimicking naturally occurring nucleosides to create error catastrophe during virus replication in the host. Therefore, due to its random nature, we would expect this compound to act against all variants of the virus in a similar manner as it does not target any specific region of the virus. Dose response assays were performed by infecting hACE2-A549 and Calu-3 cells at an MOI of 0.1 in media alone and in media containing 0.01, 0.1, 1, and 10μM NHC. After 72 hours, supernatants were removed, and viral titres determined by plaque assay. The IC50 was determined using a 4-parameter non-linear regression with GraphPad Prism 9.5. IC50 values were similar for each variant in hACE2-A549 cells of 0.1μM, while there was more variation observed in Calu-3 cells, ranging from 0.11 to 0.38μM in different variants. T-tests revealed that there was no significant difference between the IC50 for each VOC in either cell line (p>0.05) and an extra sum-of-squares F-test revealed that there was no significant difference between the best-fit curves for hACE2-A549 cells (p=0.99) and Calu-3 cells (p=0.16) (Table 1, Figure 2). Interestingly Delta had a slightly elevated IC50 compared to the other VOCs (though not statistically significant), which has been noted before in a Roborovski dwarf hamster model (Lieber et al., 2022).

**Table 1.**
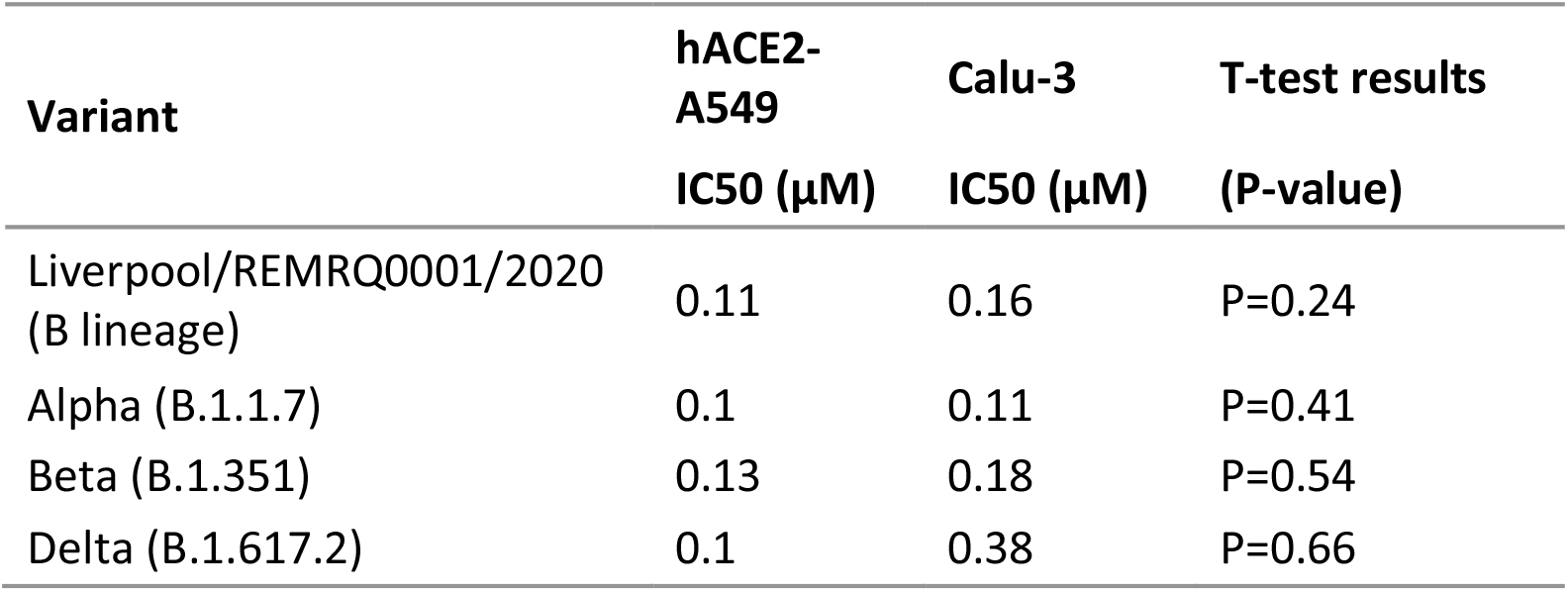
IC50 values of NHC against variants in human lung cell models. Inhibitory activity of NHC against an early variant of SARS-CoV-2 (Liverpool/REMRQ0001/2020) and Alpha (B.1.1.7), Beta (B.1.351) and Delta (B.1.617.2) variants of concern (VOCs). A four-parameter non-linear regression was used to calculate the IC50 for each variant (n=4) and T-tests used to compare the IC50 for each VOC across cell-lines.

| Variant | hACE2-A549<br>IC50 (μM) | Calu-3<br>IC50 (μM) | T-test results<br>(P-value) |
| --- | --- | --- | --- |
| Liverpool/REMRQ0001/2020<br>(B lineage) | 0.11 | 0.16 | P=0.24 |
| Alpha (B.1.1.7) | 0.1 | 0.11 | P=0.41 |
| Beta (B.1.351) | 0.13 | 0.18 | P=0.54 |
| Delta (B.1.617.2) | 0.1 | 0.38 | P=0.66 |

**Figure 2.**
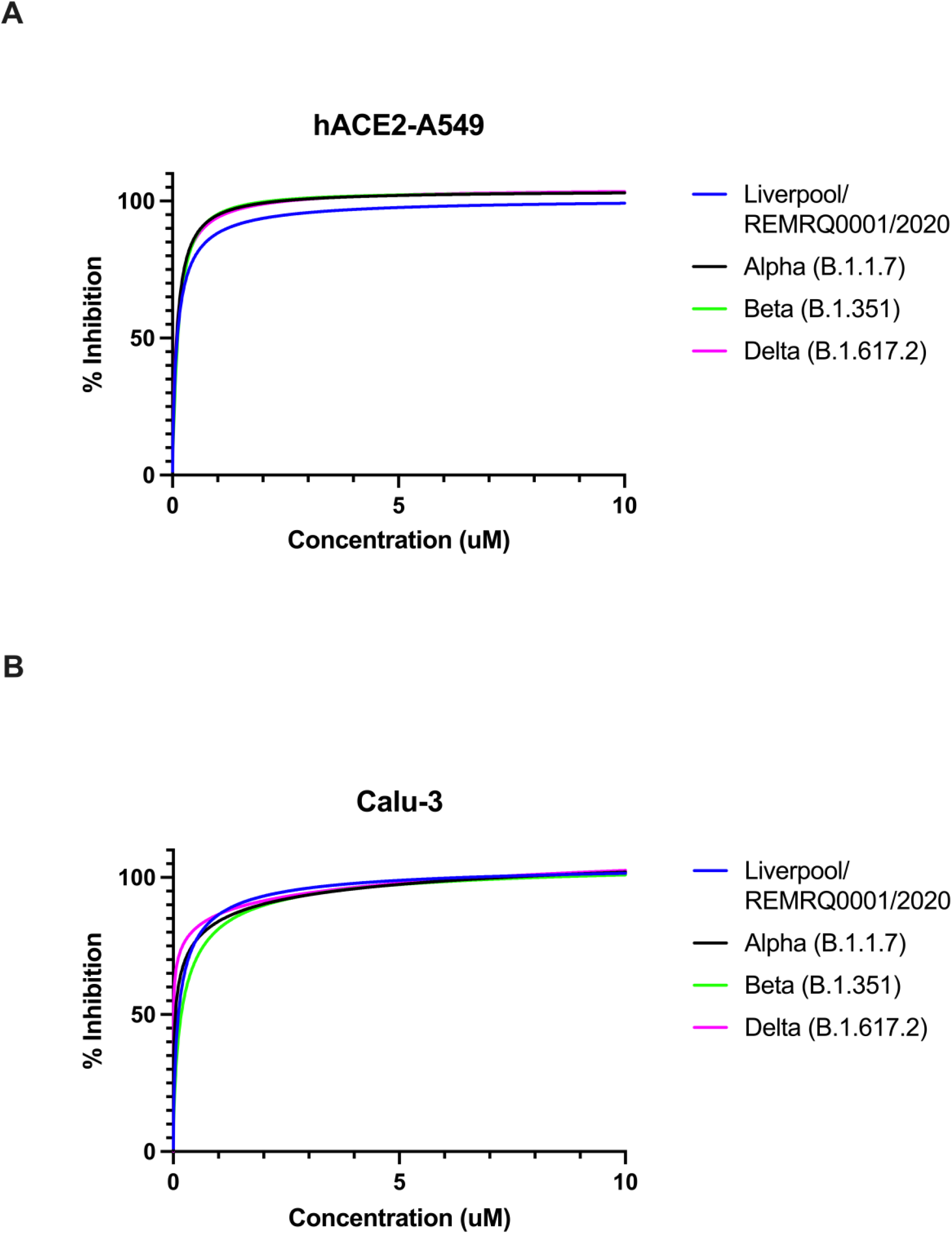
NHC is similarly active against both an early variant and VOCs in A) hACE2-A549 cells and B) Calu-3 cells. Inhibitory activity of NHC against an early variant of SARS-CoV-2 (Liverpool/REMRQ001/2020) and Alpha (B.1.1.7), Beta (B.1.351) and Delta (B.1.617.2) variants of concern (VOCs). A non-linear regression was used to plot the dose response curves and calculate the IC50 for each variant (n=4).

### 3.2 The ability of NHC to inhibit the growth of SARS-CoV-2 VOCs is dependent on timing of administration

To determine the effective treatment window, hACE2-A549 cells were both pre-treated with different concentrations of NHC two hours prior to infection and treated at two, four, 24- and 48-hours post-infection and compared to concurrent treatment/infection. Dose-response assays were performed on supernatants collected 72 hours post-infection. Viral titres remained similar if treatment was given prior to, or at the same time as infection and at time points of two, four- and 24-hours post-infection. However, if treatment was given at 48 hours post-infection, the drug was not as effective for all variants tested with inhibition never reaching 100% (Figure 3). An extra sum-of squares F-test was performed for each variant and demonstrated that the curves differed significantly for Liverpool (P=0.0018), Alpha (P<0.001), Beta (P=0.0008) but a p-value could not be calculated for Delta.

**Figure 3.**
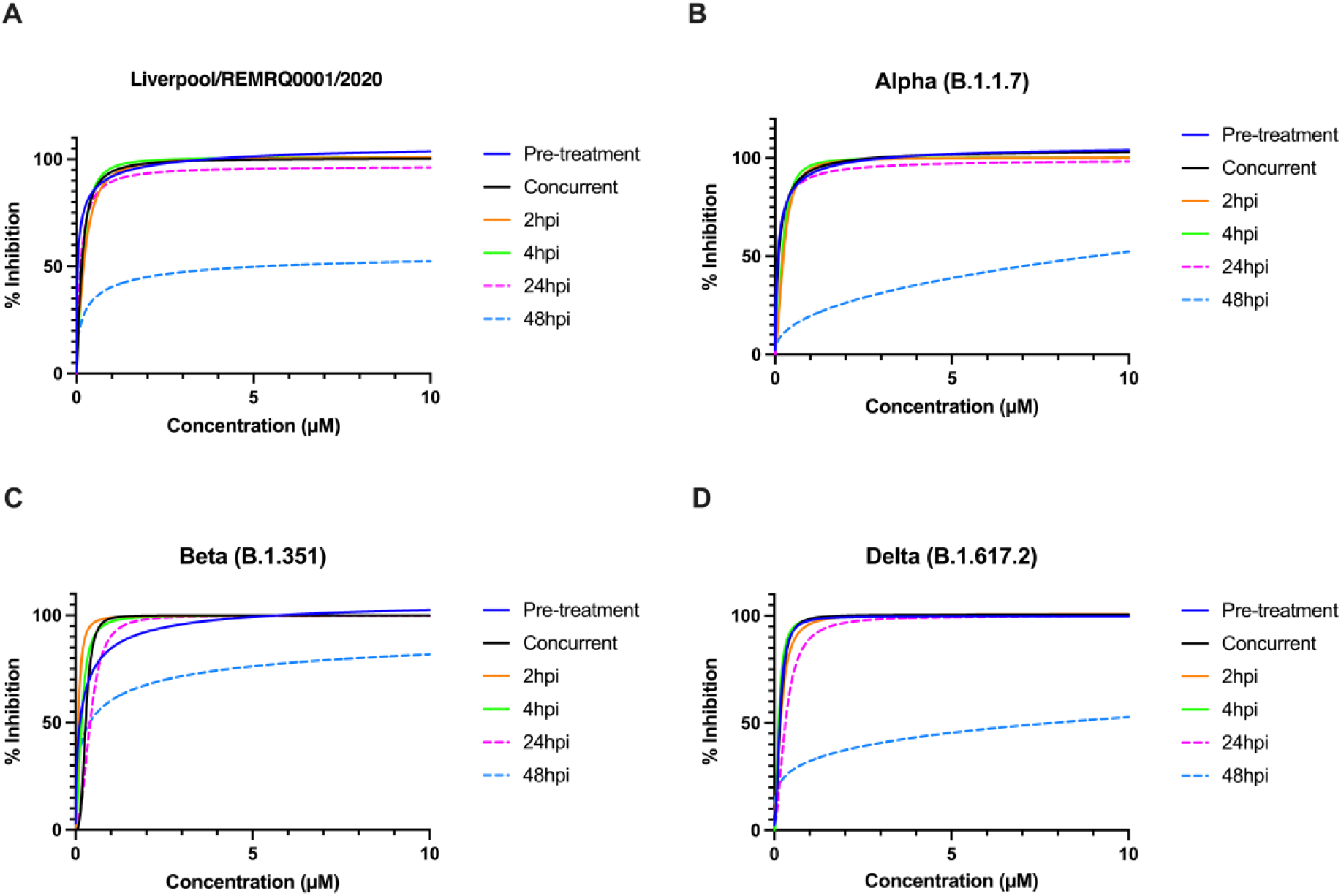
Effect of pre-treatment, concurrent treatment and treatment at two, four, 24 and 48 hours post-infection on inhibition of viral growth in hACE2-A549 cells. Dose response assays were performed on supernatants collected 72 hours post-infection for (A) the Liverpool variant, (B) Alpha (B.1.1.7), (C) Beta (B.1.351) or (D) Delta (B.1.617.2) variants. A non-linear regression was used to plot the dose response curves for each variant (n=3).

These results have been borne out by others using a hollow-fibre injection model (Brown et al., 2022) and in animal models (Abdelnabi et al., 2021; Rosenke et al., 2021). Virological and pathological analysis in patients who died of severe COVID-19 suggested that by the time immunopathology is recognised, giving an anti-viral in the expectation that this would reduce symptoms associated with COVID-19 is too late (Dorward et al., 2021). While the large clinical studies on patients receiving Molnupiravir claim patients were given the drug within five days of symptoms, few have directly measured the effect of delays in treatment post-symptom onset. A weak relationship (not statistically significant) between disease onset time and administration of Molnupiravir was identified in a single centre observational study in Poland. A delay in Molnupiravir administration was associated with an increase in the likelihood of hospitalisation (Czarnecka et al., 2022). Furthermore a recent Italian study in patients > 80 years old found that the greater the number of days from the onset of symptoms to administration of therapy, the higher the likelihood was of hospitalisation or death (Bruno et al., 2022). These results indicate that the relative muted effects of Molnupiravir given to patients in real world settings may be the result of delayed administration of the drug.

### 3.3 A fixed dose exposure to the IC50 of NHC did not induce resistance after 20 passages for Beta, but results were inconclusive for the Liverpool variant

Anti-viral drugs are being used to treat patients with persistent infection, particularly in patients who have compromised immunity. In this setting, Molnupiravir may be given over extended periods of time. Therefore, to determine if drug-induced adaptations to NHC were likely to emerge after prolonged exposure, Calu-3 cells were infected in either the presence of a sub-optimal dose of NHC (0.1μM) or in media alone, with two variants, the Liverpool/REMRQ0001/2020 (B lineage), and the Beta VOC (B.1.351). These viruses were passaged 20 times and supernatants were titred and analysed by NHC dose response assays every five passages in order to determine if the IC50 had changed.

For the Liverpool variant, logistic regression curves for each passage were plotted. An extra sum-of squares F-test suggested that the curves differed (p=0.019) (Figure 4a). The percent inhibition values are plotted in Figure 4b with the mean +/-SEM. To explore this further, the IC50 values for each replicate were plotted and an ordinary one-way ANOVA with post-hoc Tukey test was performed to determine if there was any significant change over the course of passage. This found no significant differences in the IC50 values between any pairwise combination (p=0.55) (Figure 4c).

**Figure 4.**
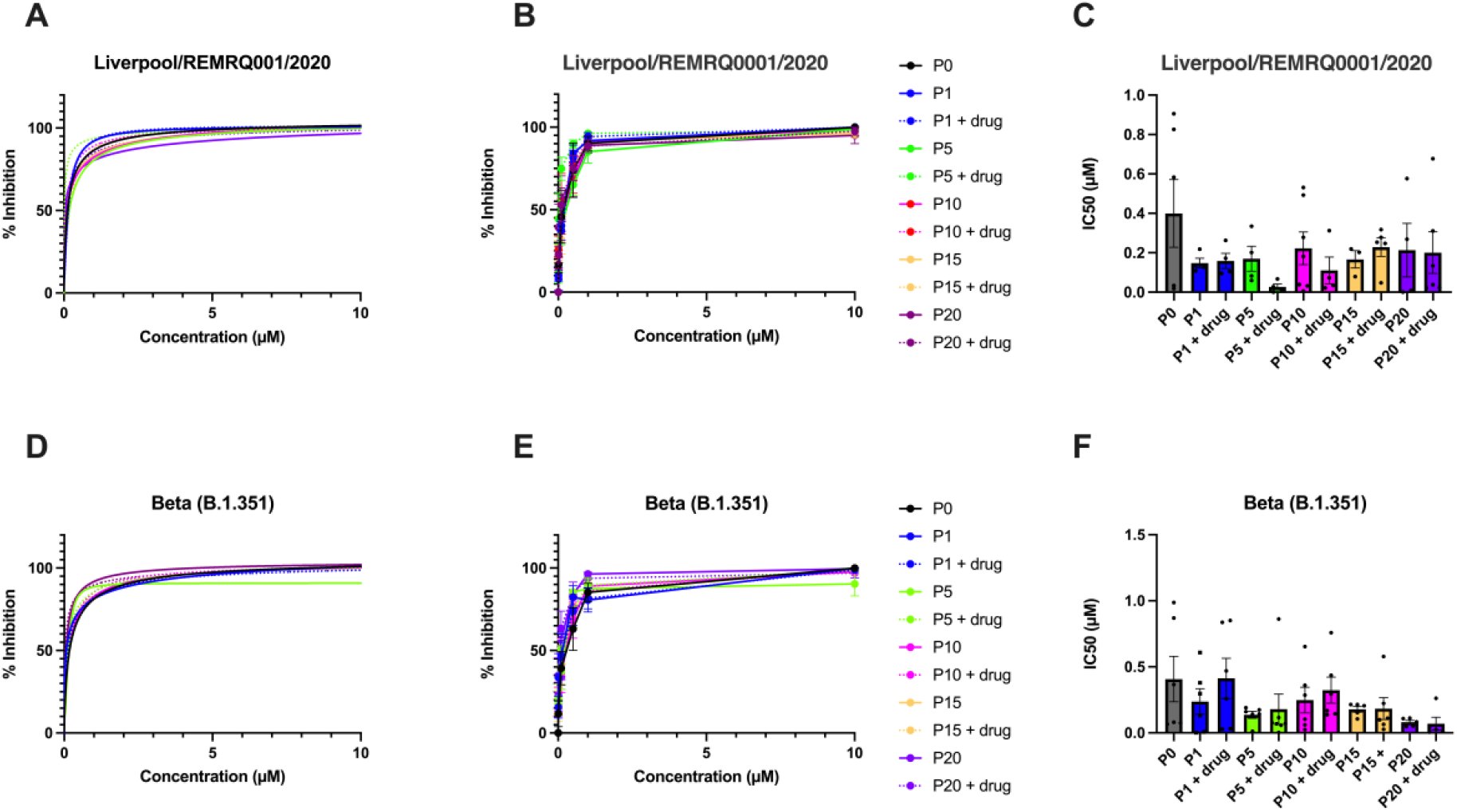
A fixed dose exposure to the IC50 of NHC did not induce resistance after 20 passages for Beta, but results were inconclusive for the Liverpool variant. (A) Logistic regression curves for the Liverpool variant. (B) Percentage inhibition values plotted for each passage (mean ± SEM) for the Liverpool variant. (C) IC50 values plotted for each passage (mean ± SEM) for the Liverpool variant (n>/=3). (D) Logistic regression curves for the Beta VOC. (E) Percentage inhibition values plotted for each passage (mean ± SEM) for the Beta VOC. (F) IC50 values plotted for each passage (mean ± SEM) for the Beta VOC (n>/=5).

Dose response results for the Beta variant were analysed by logistic regression and likewise plotted with the extra sum-of-squares F-test but the result demonstrated no difference between the curves (p=0.66) (Figure 4d). The percent inhibition values are plotted in Figure 4e with the mean +/-SEM. The IC50 were similarly plotted, and the ANOVA revealed there was no significant difference between any pairwise combination (p=0.28) (Figure 4f).

While the results for the Liverpool variant were inconclusive, the Beta variant showed no signs of resistance. Although our methods differ, most studies agree that Molnupiravir has a high genetic barrier to resistance in viruses, such as VEEV (Urakova et al., 2018)Influenza, RSV (Yoon et al., 2018) or MERS (Agostini et al., 2019). However, more studies should be performed in animal models to better delineate this.

All anti-virals currently approved for use against SARS-CoV-2 have their limitations and Molnupiravir is no exception. While results suggesting that earlier administration could improve the efficacy of the drug in real-world scenarios have been confirmed by others, Brown et al (2022) have suggested increasing the dose of the drug 5-fold could overcome delays to therapy. However, tolerability in patients could be an issue and higher doses are not recommended due to concerns over host mutagenicity (Zhou et al., 2021).

### 3.4 Sequencing of serially passaged virus revealed that the hallmark of Molnupiravir treatment was detectable *in vitro*

Molnupiravir acts to cause error catastrophe through the induction of transition mutations (e.g. C -> U or G -> A changes). We hypothesised that if this were the case, that there would have been an increase in the proportion of transition to transversion mutations with treatment, compared to control. To investigate this, one sample for each condition was taken forward to measure the transition/transversion ratio in the genome of SARS-CoV-2 during serial passage (figure 5). The data indicated that repeated exposure to NHC (e.g. in passaging) had a cumulative effect on the mean transition/transversion ratio for both variants tested. At p1, the mean transition/transversion ratio for untreated Liverpool virus was 0.18, increasing to 0.37 at p20 while for NHC treated virus the ratio went from 0.2 to 0.83. For the beta variant, the mean transition/transversion ratio in untreated samples increased from 0.43 at P1 to 0.48 at P20 while NHC treated virus increased from 0.42 to 0.88.

**Figure 5.**
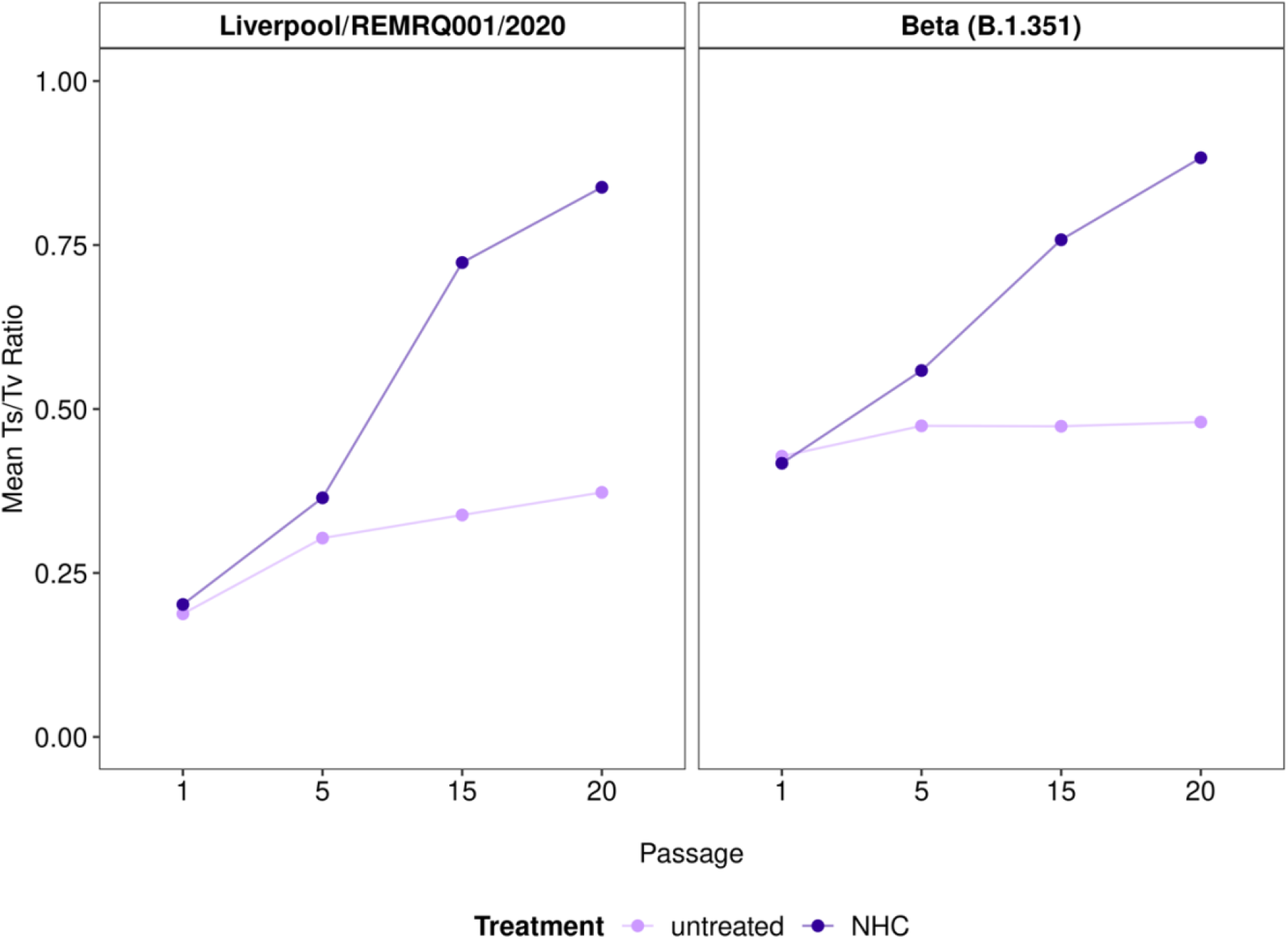
The mean transition/transversion ratio for each passaged virus. Graphs plotting the mean transition/transversion ratio (Ts/Tv) for each virus for untreated and NHC treated viruses at each passage (n=1).

### 3.5 An analysis of the non-synonymous and synonymous amino acid changes revealed that there was little difference between untreated and NHC treated viruses

To investigate the selection pressure of Molnupiravir on different viral proteins, an analysis of the predicted amino-acid changes relative to the length of each viral protein was performed. (Figure 6a and b). Transition mutations are less likely to result in non-synonymous amino-acid substitutions (Stoltzfus and Norris, 2015), so Molnupiravir would most likely cause synonymous amino acid substitutions. We did not observe any preferential regions in the genome that had an increased frequency of either synonymous or non-synonymous amino-acid substitutions in NHC treated cells compared to untreated.

**Figure 6.**
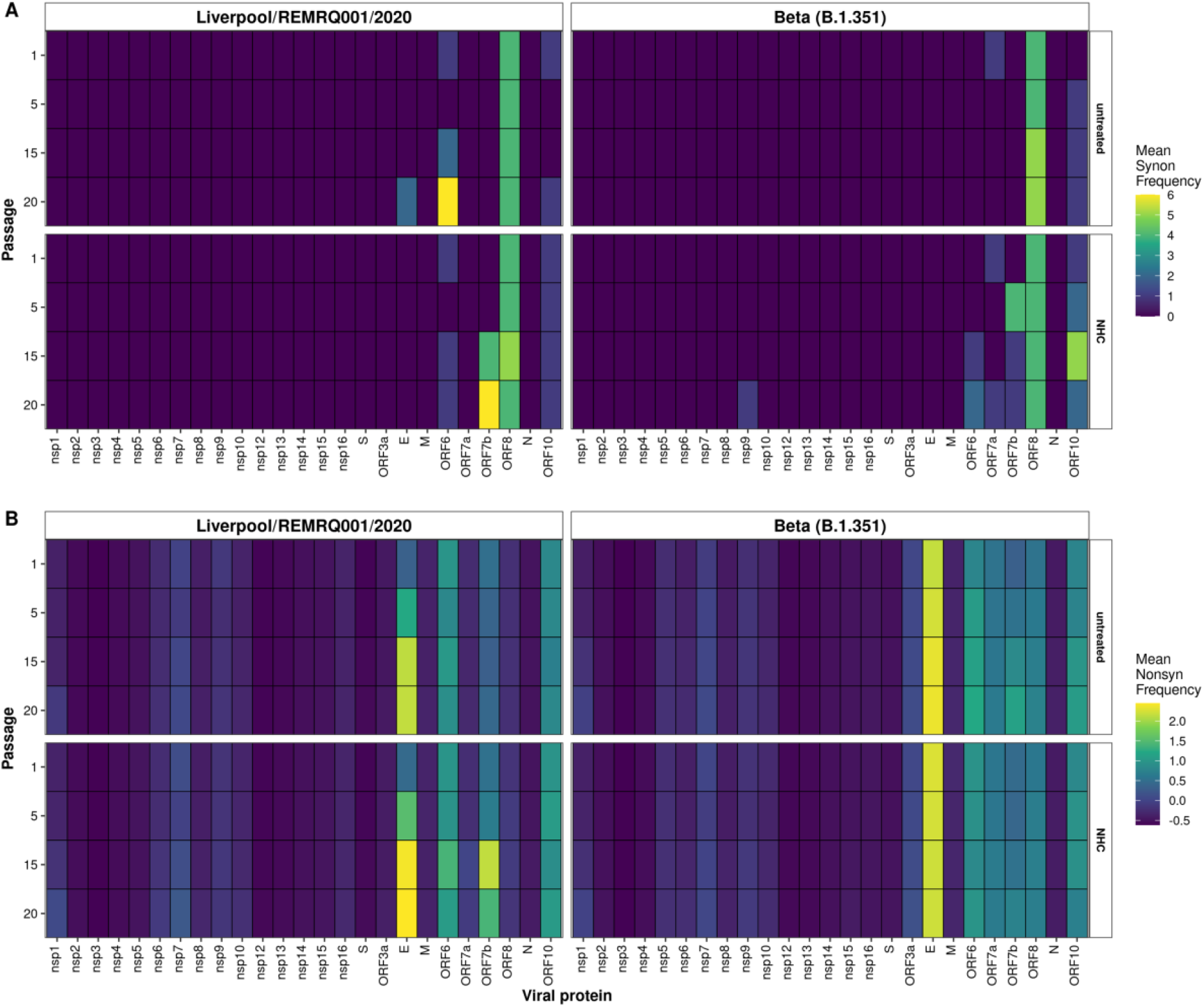
The frequency of amino acid changes relative to length of viral protein between treated and untreated viruses. Figures show the frequency (%) of amino-acid changes relative to the length of protein for each passage for both treated and untreated viruses. A) Synonymous amino acid changes. B) Non-synonymous amino acid changes. Data is missing for p10 untreated Liverpool variant (n=1).

We propose that the lack of observed resistance is due to the random nature of the drug’s mechanism of action. However, recently scientists have suggested that Molnupiravir could lead to the generation of new variants through the same mechanism (Focosi, 2022). In our work we have shown that serial passage of virus in the presence of drug resulted in a cumulative increase in the transition to transversion ratio (Figure 5). This has been noted in patients from the AGILE trial (Donovan-Banfield et al., 2022), and elsewhere it induced a number of SNPs in the viral genome which increased over time. However these SNPs were random and rarely persisted across timepoints suggesting there is less risk of the emergence of new variants (Alteri et al., 2022).

Administering the drug when patients first display symptoms may not reduce viral loads sufficiently to prevent disease. Based on the *in vitro* data, we hypothesize that Molnupiravir should be given to patients as close to exposure as possible or prophylactically. This is similar to the anti-viral philosophy underpinning the treatment of influenza infection or pre-exposure prophylaxis for HIV. In agreement with our analysis of patients treated with Molnupiravir (Donovan-Banfield et al., 2022), no changes in the predicted amino acid sequence of SARS-CoV-2 was observed between treated and untreated conditions. However, when SARS-CoV-2 was serially passaged with Molnupiravir, viable virus was recovered at each stage with an increased mutational landscape. This emphasises that whatever the treatment option in whatever patient category, if Molnupiravir is to be used, the dose given must be sufficient to eliminate virus there and then. The implication of this observation is that in persistently infected individuals, treatment should be such that virus is eliminated immediately or risk increased mutational frequency. We predict that SARS-CoV-2 genomes derived from cells that received Molnupiravir will have higher Ts/Tv ratios and at a population level these may be distinguishable during genomic surveillance.

## Acknowledgments

Virus isolates used in this study were originally derived from samples gathered under the auspices of the ISARIC Coronavirus Clinical Characterisation Consortium (ISARIC4C) and processed at the University of Liverpool. We would like to acknowledge all members of the consortium.

## Author contributions

Conceptualization: TP, SK and JAH. Methodology: TP, ID-B, HG and RP-R. Validation: TP. Formal analysis: TP, ID-B, HG, RP-R and JAH. Investigation: TP. Resources: SK and JAH. Data Curation: TP, ID-B, HG and RP-R. Writing – original draft: TP and JAH. Writing – Review and Editing: All authors. Visualisation: TP. Supervision: TF, SK and JAH. Project administration: JAH. Funding acquisition: TF and JAH.

## Bibliography

Abdelnabi, R., Foo, C.S., Kaptein, S.J.F., Zhang, X., Do, T.N.D., Langendries, L., Vangeel, L., Breuer, J., Pang, J., Williams, R., Vergote, V., Heylen, E., Leyssen, P., Dallmeier, K., Coelmont, L., Chatterjee, A.K., Mols, R., Augustijns, P., De Jonghe, S., Jochmans, D., Weynand, B., Neyts, J., 2021. The combined treatment of Molnupiravir and Favipiravir results in a potentiation of antiviral efficacy in a SARS-CoV-2 hamster infection model. EBioMedicine 72, 103595.

Agostini, M.L., Pruijssers, A.J., Chappell, J.D., Gribble, J., Lu, X., Andres, E.L., Bluemling, G.R., Lockwood, M.A., Sheahan, T.P., Sims, A.C., Natchus, M.G., Saindane, M., Kolykhalov, A.A., Painter, G.R., Baric, R.S., Denison, M.R., 2019. Small-Molecule Antiviral β-d-N(4)-Hydroxycytidine Inhibits a Proofreading-Intact Coronavirus with a High Genetic Barrier to Resistance. J Virol 93.

Alteri, C., Fox, V., Scutari, R., Burastero, G.J., Volpi, S., Faltoni, M., Fini, V., Granaglia, A., Esperti, S., Gallerani, A., Costabile, V., Fontana, B., Franceschini, E., Meschiari, M., Campana, A., Bernardi, S., Villani, A., Bernaschi, P., Russo, C., Guaraldi, G., Mussini, C., Perno, C., 2022. Genomic Evolution of Sars-Cov-2 in Molnupiravir-Treated Patients Compared to Paxlovid-Treated and Drug-Naïve Patients: A Proof-of-Concept Study. Research Square.

Bojkova, D., Widera, M., Ciesek, S., Wass, M.N., Michaelis, M., Cinatl, J., 2022. Reduced interferon antagonism but similar drug sensitivity in Omicron variant compared to Delta variant of SARS-CoV-2 isolates. Cell Research 32, 319–321.

Brown, A., N., Lang, Y., Zhou, J., Franco Evelyn, J., Hanrahan Kaley, C., Bulitta Juergen, B., Drusano George, L., 2022. Why Molnupiravir Fails in Hospitalized Patients. mBio 0, e02916–02922.

Bruno, G., Perelli, S., Giotta, M., Bartolomeo, N., De Vita, G., Buccoliero, G.B., 2022. Efficacy and safety of oral antivirals in individuals aged 80 years or older with mild-to-moderate COVID-19: preliminary report from an Italian Prescriber Center. Infez Med 30, 547–554.

Buchrieser, J., Dufloo, J., Hubert, M., Monel, B., Planas, D., Rajah, M.M., Planchais, C., Porrot, F., Guivel-Benhassine, F., Van der Werf, S., Casartelli, N., Mouquet, H., Bruel, T., Schwartz, O., 2020. Syncytia formation by SARS-CoV-2-infected cells. The EMBO Journal 39, e106267.

Butler, C.C., Hobbs, F.D.R., Gbinigie, O.A., Rahman, N.M., Hayward, G., Richards, D.B., Dorward, J., Lowe, D.M., Standing, J.F., Breuer, J., Khoo, S., Petrou, S., Hood, K., Nguyen-Van-Tam, J.S., Patel, M.G., Saville, B.R., Marion, J., Ogburn, E., Allen, J., Rutter, H., Francis, N., Thomas, N.P.B., Evans, P., Dobson, M., Madden, T.-A., Holmes, J., Harris, V., Png, M.E., Lown, M., van Hecke, O., Detry, M.A., Saunders, C.T., Fitzgerald, M., Berry, N.S., Mwandigha, L., Galal, U., Mort, S., Jani, B.D., Hart, N.D., Ahmed, H., Butler, D., McKenna, M., Chalk, J., Lavallee, L., Hadley, E., Cureton, L., Benysek, M., Andersson, M., Coates, M., Barrett, S., Bateman, C., Davies, J.C., Raymundo-Wood, I., Ustianowski, A., Carson-Stevens, A., Yu, L.-M., Little, P., Agyeman, A.A., Ahmed, T., Allcock, D., Beltran-Martinez, A., Benedict, O.E., Bird, N., Brennan, L., Brown, J., Burns, G., Butler, M., Cheng, Z., Danson, R., de Kare-Silver, N., Dhasmana, D., Dickson, J., Engamba, S., Fisher, S., Fox, R., Frost, E., Gaunt, R., Ghosh, S., Gilkar, I., Goodman, A., Granier, S., Howell, A., Hussain, I., Hutchinson, S., Imlach, M., Irving, G., Jacobsen, N., Kennard, J., Khan, U., Knox, K., Krasucki, C., Law, T., Lee, R., Lester, N., Lewis, D., Lunn, J., Mackintosh, C.I., Mathukia, M., Moore, P., Morton, S., Murphy, D., Nally, R., Ndukauba, C., Ogundapo, O., Okeke, H., Patel, A., Patel, K., Penfold, R., Poonian, S., Popoola, O., Pora, A., Prasad, V., Prasad, R., Razzaq, O., Richardson, S., Royal, S., Safa, A., Sehdev, S., Sevenoaks, T., Shah, D., Sheikh, A., Short, V., Sidhu, B.S., Singh, I., Soni, Y., Thalasselis, C., Wilson, P., Wingfield, D., Wong, M., Woodall, M.N.J., Wooding, N., Woods, S., Yong, J., Yongblah, F., Zafar, A., 2022. Molnupiravir plus usual care versus usual care alone as early treatment for adults with COVID-19 at increased risk of adverse outcomes (PANORAMIC): an open-label, platform-adaptive randomised controlled trial. The Lancet.

Coolen, J.P.M., Wolters, F., Tostmann, A., van Groningen, L.F.J., Bleeker-Rovers, C.P., Tan, E., van der Geest-Blankert, N., Hautvast, J.L.A., Hopman, J., Wertheim, H.F.L., Rahamat-Langendoen, J.C., Storch, M., Melchers, W.J.G., 2021. SARS-CoV-2 whole-genome sequencing using reverse complement PCR: For easy, fast and accurate outbreak and variant analysis. J Clin Virol 144, 104993.

Cox, M., Peacock, T.P., Harvey, W.T., Hughes, J., Wright, D.W., Willett, B.J., Thomson, E., Gupta, R.K., Peacock, S.J., Robertson, D.L., Carabelli, A.M., Consortium, C.-G.U., 2022. SARS-CoV-2 variant evasion of monoclonal antibodies based on in vitro studies. Nature Reviews Microbiology.

Cox, R.M., Wolf, J.D., Plemper, R.K., 2021. Therapeutically administered ribonucleoside analogue MK-4482/EIDD-2801 blocks SARS-CoV-2 transmission in ferrets. Nat Microbiol 6, 11–18.

Czarnecka, K., Czarnecka, P., Tronina, O., Durlik, M., 2022. Molnupiravir Outpatient Treatment for Adults with COVID-19 in a Real-World Setting—A Single Center Experience, Journal of Clinical Medicine.

Donovan-Banfield, I.a., Penrice-Randal, R., Goldswain, H., Rzeszutek, A.M., Pilgrim, J., Bullock, K., Saunders, G., Northey, J., Dong, X., Ryan, Y., Reynolds, H., Tetlow, M., Walker, L.E., FitzGerald, R., Hale, C., Lyon, R., Woods, C., Ahmad, S., Hadjiyiannakis, D., Periselneris, J., Knox, E., Middleton, C., Lavelle-Langham, L., Shaw, V., Greenhalf, W., Edwards, T., Lalloo, D.G., Edwards, C.J., Darby, A.C., Carroll, M.W., Griffiths, G., Khoo, S.H., Hiscox, J.A., Fletcher, T., 2022. Characterisation of SARS-CoV-2 genomic variation in response to Molnupiravir treatment in the AGILE Phase IIa clinical trial. Nature Communications 13, 7284.

Dorward, D.A., Russell, C.D., Um, I.H., Elshani, M., Armstrong, S.D., Penrice-Randal, R., Millar, T., Lerpiniere, C.E.B., Tagliavini, G., Hartley, C.S., Randle, N.P., Gachanja, N.N., Potey, P.M.D., Dong, X., Anderson, A.M., Campbell, V.L., Duguid, A.J., Al Qsous, W., BouHaidar, R., Baillie, J.K., Dhaliwal, K., Wallace, W.A., Bellamy, C.O.C., Prost, S., Smith, C., Hiscox, J.A., Harrison, D.J., Lucas, C.D., 2021. Tissue-Specific Immunopathology in Fatal COVID-19. Am J Respir Crit Care Med 203, 192–201.

Fischer, W.A., Eron, J.J., Holman, W., Cohen, M.S., Fang, L., Szewczyk, L.J., Sheahan, T.P., Baric, R., Mollan, K.R., Wolfe, C.R., Duke, E.R., Azizad, M.M., Borroto-Esoda, K., Wohl, D.A., Coombs, R.W., Loftis, A.J., Alabanza, P., Lipansky, F., Painter, W., 2021. A Phase 2a clinical trial of Molnupiravir in patients with COVID-19 shows accelerated SARS-CoV-2 RNA clearance and elimination of infectious virus. Science Translational Medicine 0, eabl7430.

Focosi, D., 2022. Molnupiravir: From Hope to Epic Fail?, Viruses.

Haddad, F., Dokmak, G., Karaman, R., 2022. A Comprehensive Review on the Efficacy of Several Pharmacologic Agents for the Treatment of COVID-19, Life.

Jayk Bernal, A., Gomes da Silva, M.M., Musungaie, D.B., Kovalchuk, E., Gonzalez, A., Delos Reyes, V., Martín-Quirós, A., Caraco, Y., Williams-Diaz, A., Brown, M.L., Du, J., Pedley, A., Assaid, C., Strizki, J., Grobler, J.A., Shamsuddin, H.H., Tipping, R., Wan, H., Paschke, A., Butterton, J.R., Johnson, M.G., De Anda, C., 2022. Molnupiravir for Oral Treatment of Covid-19 in Nonhospitalized Patients. New England Journal of Medicine.

Jochmans, D., Liu, C., Donckers, K., Stoycheva, A., Boland, S., Stevens, S.K., De Vita, C., Vanmechelen, B., Maes, P., Trüeb, B., Ebert, N., Thiel, V., De Jonghe, S., Vangeel, L., Bardiot, D., Jekle, A., Blatt, L.M., Beigelman, L., Symons, J.A., Raboisson, P., Chaltin, P., Marchand, A., Neyts, J., Deval, J., Vandyck, K., 2023. The substitutions L50F, E166A and L167F in SARS-CoV-2 3CLpro are selected by a protease inhibitor in vitro; and confer resistance to nirmatrelvir. mBio e02815–22.

Johnson, M.G., Strizki, J.M., Brown, M.L., Wan, H., Shamsuddin, H.H., Ramgopal, M., Florescu, D.F., Delobel, P., Khaertynova, I., Flores, J.F., Fouche, L.F., Chang, S.-C., Williams-Diaz, A., Du, J., Grobler, J.A., Paschke, A., De Anda, C., 2023. Molnupiravir for the treatment of COVID-19 in immunocompromised participants: efficacy, safety, and virology results from the phase 3 randomized, placebo-controlled MOVe-OUT trial. Infection.

Kabinger, F., Stiller, C., Schmitzová, J., Dienemann, C., Kokic, G., Hillen, H.S., Höbartner, C., Cramer, P., 2021. Mechanism of Molnupiravir-induced SARS-CoV-2 mutagenesis. Nature Structural & Molecular Biology.

Kearns, P., Siebert, S., Willicombe, m., Gaskell, C., Kirkham, A., Pirrie, S., Bowden, S., Magwaro, S., Hughes, A., Lim, Z., Dimitriadis, S., Murray, S.M., Marjot, T., Win, Z., Irwin, S.L., Meacham, G., Richter, A.G., Kelleher, P., Satsangi, J., Miller, P., Rea, D., Cook, G., Turtle, L., Klenerman, P., Dunachie, S., Basu, N., de Silva, T.I., Thomas, D., Barnes, E., Goodyear, C.S., McInnes, I., 2021. Examining the Immunological Effects of COVID-19 Vaccination in Patients with Conditions Potentially Leading to Diminished Immune Response Capacity – The OCTAVE Trial., Preprintes with The Lancet.

Khoo, S.H., FitzGerald, R., Saunders, G., Middleton, C., Ahmad, S., Edwards, C.J., Hadjiyiannakis, D., Walker, L., Lyon, R., Shaw, V., Mozgunov, P., Periselneris, J., Woods, C., Bullock, K., Hale, C., Reynolds, H., Downs, N., Ewings, S., Buadi, A., Cameron, D., Edwards, T., Knox, E., Donovan-Banfield, I.a., Greenhalf, W., Chiong, J., Lavelle-Langham, L., Jacobs, M., Northey, J., Painter, W., Holman, W., Lalloo, D.G., Tetlow, M., Hiscox, J.A., Jaki, T., Fletcher, T., Griffiths, G., Paton, N., Hayden, F., Darbyshire, J., Lucas, A., Lorch, U., Freedman, A., Knight, R., Julious, S., Byrne, R., Cubas Atienzar, A., Jones, J., Williams, C., Song, A., Dixon, J., Alexandersson, A., Hatchard, P., Tilt, E., Titman, A., Doce Carracedo, A., Chandran Gorner, V., Davies, A., Woodhouse, L., Carlucci, N., Okenyi, E., Bula, M., Dodd, K., Gibney, J., Dry, L., Rashid Gardner, Z., Sammour, A., Cole, C., Rowland, T., Tsakiroglu, M., Yip, V., Osanlou, R., Stewart, A., Parker, B., Turgut, T., Ahmed, A., Starkey, K., Subin, S., Stockdale, J., Herring, L., Baker, J., Oliver, A., Pacurar, M., Owens, D., Munro, A., Babbage, G., Faust, S., Harvey, M., Pratt, D., Nagra, D., Vyas, A., 2022. Molnupiravir versus placebo in unvaccinated and vaccinated patients with early SARS-CoV-2 infection in the UK (AGILE CST-2): a randomised, placebo-controlled, double-blind, phase 2 trial. The Lancet Infectious Diseases.

Kokic, G., Hillen, H.S., Tegunov, D., Dienemann, C., Seitz, F., Schmitzova, J., Farnung, L., Siewert, A., Höbartner, C., Cramer, P., 2021. Mechanism of SARS-CoV-2 polymerase stalling by remdesivir. Nature Communications 12, 279.

Lee, J., Lee, J., Kim, H.J., Ko, M., Jee, Y., Kim, S., 2021. TMPRSS2 and RNA-dependent RNA polymerase are effective targets of therapeutic intervention for treatment of COVID-19 caused by SARS-CoV-2 variants (B.1.1.7 and B.1.351). Microbiol Spectr 9, e0047221.

Lieber, C.M., Cox, R.M., Sourimant, J., Wolf, J.D., Juergens, K., Phung, Q., Saindane, M.T., Smith, M.K., Sticher, Z.M., Kalykhalov, A.A., Natchus, M.G., Painter, G.R., Sakamoto, K., Greninger, A.L., Plemper, R.K., 2022. SARS-CoV-2 VOC type and biological sex affect Molnupiravir efficacy in severe COVID-19 dwarf hamster model. Nature Communications 13, 4416.

Moghadasi, S.A., Heilmann, E., Moraes, S.N., Kearns, F.L., von Laer, D., Amaro, R.E., Harris, R.S., 2022. Transmissible SARS-CoV-2 variants with resistance to clinical protease inhibitors. bioRxiv.

O’Toole, Á., Scher, E., Underwood, A., Jackson, B., Hill, V., McCrone, J.T., Colquhoun, R., Ruis, C., Abu-Dahab, K., Taylor, B., Yeats, C., du Plessis, L., Maloney, D., Medd, N., Attwood, S.W., Aanensen, D.M., Holmes, E.C., Pybus, O.G., Rambaut, A., 2021. Assignment of epidemiological lineages in an emerging pandemic using the pangolin tool. Virus Evolution 7.

Rosenke, K., Hansen, F., Schwarz, B., Feldmann, F., Haddock, E., Rosenke, R., Barbian, K., Meade-White, K., Okumura, A., Leventhal, S., Hawman, D.W., Ricotta, E., Bosio, C.M., Martens, C., Saturday, G., Feldmann, H., Jarvis, M.A., 2021. Orally delivered MK-4482 inhibits SARS-CoV-2 replication in the Syrian hamster model. Nature Communications 12, 2295.

Sheahan, T.P., Sims, A.C., Zhou, S., Graham, R.L., Pruijssers, A.J., Agostini, M.L., Leist, S.R., Schäfer, A., Dinnon, K.H., Stevens, L.J., Chappell, J.D., Lu, X., Hughes, T.M., George, A.S., Hill, C.S., Montgomery, S.A., Brown, A.J., Bluemling, G.R., Natchus, M.G., Saindane, M., Kolykhalov, A.A., Painter, G., Harcourt, J., Tamin, A., Thornburg, N.J., Swanstrom, R., Denison, M.R., Baric, R.S., 2020. An orally bioavailable broad-spectrum antiviral inhibits SARS-CoV-2 in human airway epithelial cell cultures and multiple coronaviruses in mice. Science Translational Medicine 12, eabb5883.

Stegmann, K.M., Dickmanns, A., Heinen, N., Groß, U., Görlich, D., Pfaender, S., Dobbelstein, M., 2021. N4-hydroxycytidine and inhibitors of dihydroorotate dehydrogenase synergistically suppress SARS-CoV-2 replication. bioRxiv, 2021.2006.2028.450163.

Stoltzfus, A., Norris, R., 2015. On the Causes of Evolutionary Transition:Transversion Bias. Molecular biology and evolution 33.

Szemiel, A.M., Merits, A., Orton, R.J., MacLean, O.A., Pinto, R.M., Wickenhagen, A., Lieber, G., Turnbull, M.L., Wang, S., Furnon, W., Suarez, N.M., Mair, D., da Silva Filipe, A., Willett, B.J., Wilson, S.J., Patel, A.H., Thomson, E.C., Palmarini, M., Kohl, A., Stewart, M.E., 2021. In vitro selection of Remdesivir resistance suggests evolutionary predictability of SARS-CoV-2. PLOS Pathogens 17, e1009929.

Urakova, N., Kuznetsova, V., Crossman, D.K., Sokratian, A., Guthrie, D.B., Kolykhalov, A.A., Lockwood, M.A., Natchus, M.G., Crowley, M.R., Painter, G.R., Frolova, E.I., Frolov, I., 2018. β-d-N(4)-Hydroxycytidine Is a Potent Anti-alphavirus Compound That Induces a High Level of Mutations in the Viral Genome. J Virol 92.

Wahl, A., Gralinski, L.E., Johnson, C.E., Yao, W., Kovarova, M., Dinnon, K.H., 3rd, Liu, H., Madden, V.J., Krzystek, H.M., De, C., White, K.K., Gully, K., Schäfer, A., Zaman, T., Leist, S.R., Grant, P.O., Bluemling, G.R., Kolykhalov, A.A., Natchus, M.G., Askin, F.B., Painter, G., Browne, E.P., Jones, C.D., Pickles, R.J., Baric, R.S., Garcia, J.V., 2021. SARS-CoV-2 infection is effectively treated and prevented by EIDD-2801. Nature 591, 451–457.

Yoon, J.J., Toots, M., Lee, S., Lee, M.E., Ludeke, B., Luczo, J.M., Ganti, K., Cox, R.M., Sticher, Z.M., Edpuganti, V., Mitchell, D.G., Lockwood, M.A., Kolykhalov, A.A., Greninger, A.L., Moore, M.L., Painter, G.R., Lowen, A.C., Tompkins, S.M., Fearns, R., Natchus, M.G., Plemper, R.K., 2018. Orally Efficacious Broad-Spectrum Ribonucleoside Analog Inhibitor of Influenza and Respiratory Syncytial Viruses. Antimicrob Agents Chemother 62.

Zhou, S., Hill, C.S., Sarkar, S., Tse, L.V., Woodburn, B.M.D., Schinazi, R.F., Sheahan, T.P., Baric, R.S., Heise, M.T., Swanstrom, R., 2021. β-d-N4-hydroxycytidine Inhibits SARS-CoV-2 Through Lethal Mutagenesis But Is Also Mutagenic To Mammalian Cells. The Journal of Infectious Diseases 224, 415–419.

Zhou, Y., Gammeltoft, K.A., Ryberg, L.A., Pham, L.V., Fahnøe, U., Binderup, A., Hernandez, C.R.D., Offersgaard, A., Fernandez-Antunez, C., Peters, G.H.J., Ramirez, S., Bukh, J., Gottwein, J.M., 2022. Nirmatrelvir Resistant SARS-CoV-2 Variants with High Fitness in Vitro. Science Advances 8.

