## Supplementary figures and images for "Antiviral activity of Molnupiravir precursor NHC against SARS-CoV-2 Variants of Concern (VOCs) and implications for the therapeutic window and resistance"

### Supplementary Figure 1a and b

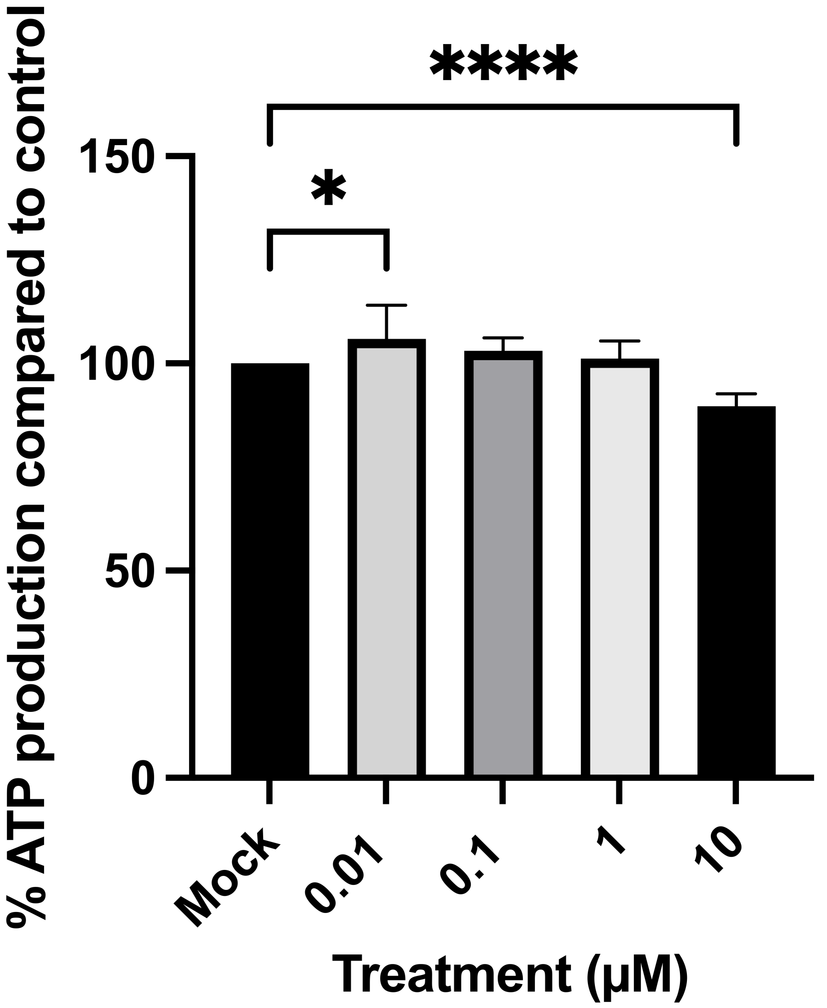

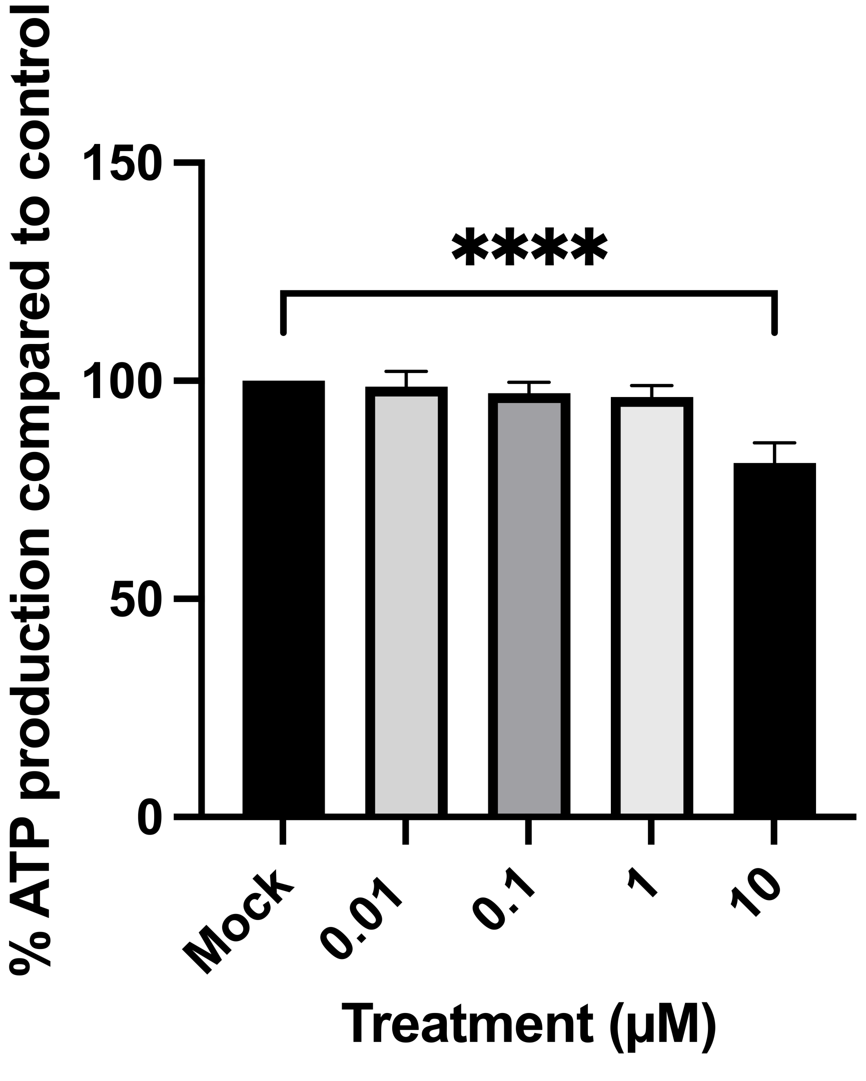
